# Is whole-brain functional connectivity a neuromarker of sustained attention? Comment on Rosenberg & al. (2016)

**DOI:** 10.1101/216697

**Authors:** Simon Bang Kristensen, Kristian Sandberg

## Abstract

Identification of neuromarkers accurately predicting cognitive characteristics from a single, standardised MRI scan could be tremendously useful in basic psychology and clinical practise. In a recent article, Rosenberg et al. (Nat Neurosci, 19, 165-171, 2016) argue that whole-brain functional network strength is a broadly applicable neuromarker of sustained attention. They claim that this marker accurately predicts performance from task related as well as resting state activity. Here, we discuss the applicability and generalizability in the context of three methodological concerns: Simulations show that the statistical methods for the 1) initial validation analyses as well as 2) internal validation using leave-one-out cross validation, are biased towards significance; 3) simple and complex models are compared suboptimally. Overall, we find that the article of Rosenberg et al. provides sufficient proof that network strength is associated with attentional capacity that it is not possible to say to which extent, and for this reason we argue that it cannot be concluded that the network is a broadly applicable neuromarker.

---

Being able to accurately predict cognitive characteristics of an individual based on a single, standardised brain scan could be tremendously useful in basic neuroscientific research and in clinical practise. The topic is currently receiving significant scientific interest with *NeuroImage* dedicating a special issue to it (Calhoun et al., 2017) and *Trends in Cognitive Sciences* publishing a Feature Review specifically on examining individual differences using functional magnetic resonance imaging (fMRI) (Dubois and Adolphs, 2016).

In a young field such as this, new methodologies for analyses are proposed, and the limits of what can be accomplished are tested. In a recent article, Rosenberg and colleagues (2016) did exactly this within the field of sustained attention. They created network models, referred to as Sustained Attention Network (SAN) models, from fMRI data and, using these, they claimed to demonstrate “that whole-brain functional network strength provides a broadly applicable neuromarker of sustained attention” (p. 165). The claim was backed by analyses showing that the marker accurately predicted not only task performance from task-related activity, but also from resting state activity, and the identified network predicted symptoms of attention deficit hyperactivity disorder (ADHD) from resting state activity in a separate sample.

While we find that the authors convincingly demonstrate that functional connectivity is associated with (sustained) attentional capacity, we would like to raise some methodological concerns, which leave us in doubt about how to interpret the created network models and, more generally, whether the article can be taken as evidence that whole-brain functional connectivity is indeed a broadly applicable neuromarker of sustained attention, or simply one of presumably many moderately predictive neural characteristics.

Our points generally fall within two overall domains, 1) specific methodological concerns and 2) a broader discussion of predictive modelling. We begin by summarising the main analysis applied in Rosenberg et al. and we present three specific methodological concerns: 1a) the network validation analysis and 1b) the correlations obtained from cross validation constitute biased tests, and 1c) complex and simple models were not compared in a principled manner. Before presenting our discussion we briefly comment on two valid analyses reported in Rosenberg et al., and what might be concluded from them. We then present a broader discussion of predictive modelling in the context of neuromarkers where we begin by 2a) briefly discussing the term ‘neuromarker’ as it appears in central claims of Rosenberg et al. Afterwards we discuss 2b) how to measure predictive power. We present these points within the frame of the central claims that the neuromarker is robust (p. 165, p. 169), predictive (pp. 165-70), and broadly applicable (p. 165), and in the final part of the comment we generalise our discussion and make recommendations for future studies predicting individual differences and identifying neuromarkers.

## 2. Summary of the general analysis method of Rosenberg et al. (2016)

The main data of the Rosenberg et al. study was the fMRI BOLD signal recorded during a sustained attention task and during resting, and the general method for model building was as follows: The brain of each participant was divided into 268 distinct nodes, and the BOLD signal of all constituent voxels was averaged within each node, creating 268 mean BOLD time courses. To examine the connectivity across areas, the authors defined a correlation matrix of the 268 nodes in their atlas (a 268 × 268 matrix where only the upper/lower triangular is of interest – a total of 35778 correlations). Next, these 35778 correlation coefficients were correlated with task performance, d´, in the sustained attention task, and significant results at a threshold of p<0.01 were used in further analyses. The correlations of the remaining data were transformed into the real numbers by Fisher’s z, and the negative and positive network strengths were defined as the sum of negative/positive z-values respectively. This general procedure for obtaining network strengths was used in a series of analyses as we describe below: a network validation analysis, internal validation tests using cross-validation (here, network strengths were created and tested on different participants), and external validation tests (here, models were built and tested on different datasets).

## 3. Methodological concerns

### 3.1 Network validation analysis

To validate the use of network strength as a summary statistic, two tests were performed based on the above-mentioned preselection of data: A test of no correlation between d´ and the negative and positive network. For these tests, large *r* (0.95 and −0.97) and low *p* (1.3 × 10^−13^ and 2.4 × 10^−15^) values were obtained, and the authors concluded that “[n]etwork strength correlated with d′ across subjects, validating its use as a summary statistic” (p. 166). However, as it had already been tested that every data point used to calculate the negative and positive network strength correlated with d′ in a particular direction, the analysis becomes circular (see e.g. Kriegeskorte et al., 2009), and a significant result is not surprising. Therefore, in our view, the test cannot be used as validation.

Furthermore, a reader may be tempted to conclude from the reported test statistics (as indeed the *Nature Neuroscience News and Views* article (Smith, 2016) covering the study does), but the reported p-values are very difficult to interpret, the distribution of parameters will be off, estimates biased and confidence intervals artificially narrow, making such conclusions meaningless.

To illustrate these issues, we performed a simulation showing that significant results are obtained the majority of times that Rosenberg et al.’s network validation test is applied to random data. We created a simulated data set for which d´ was Gaussian with a mean=2 and SD=1 (similar to the values reported by Rosenberg et al.) and with 300 “edges” of independent, uniformly distributed correlation coefficients (range: -.95 to .95), but with d´ and the correlation coefficients simulated independently. 500 simulations were performed to emulate the analysis of the positive network. The p-values obtained are presented in Figure 1a and should follow the blue uniform distribution if the method were unbiased, but instead a bimodal distribution was obtained with the majority of the distribution falling under p = 0.05.

**Figure 1:**
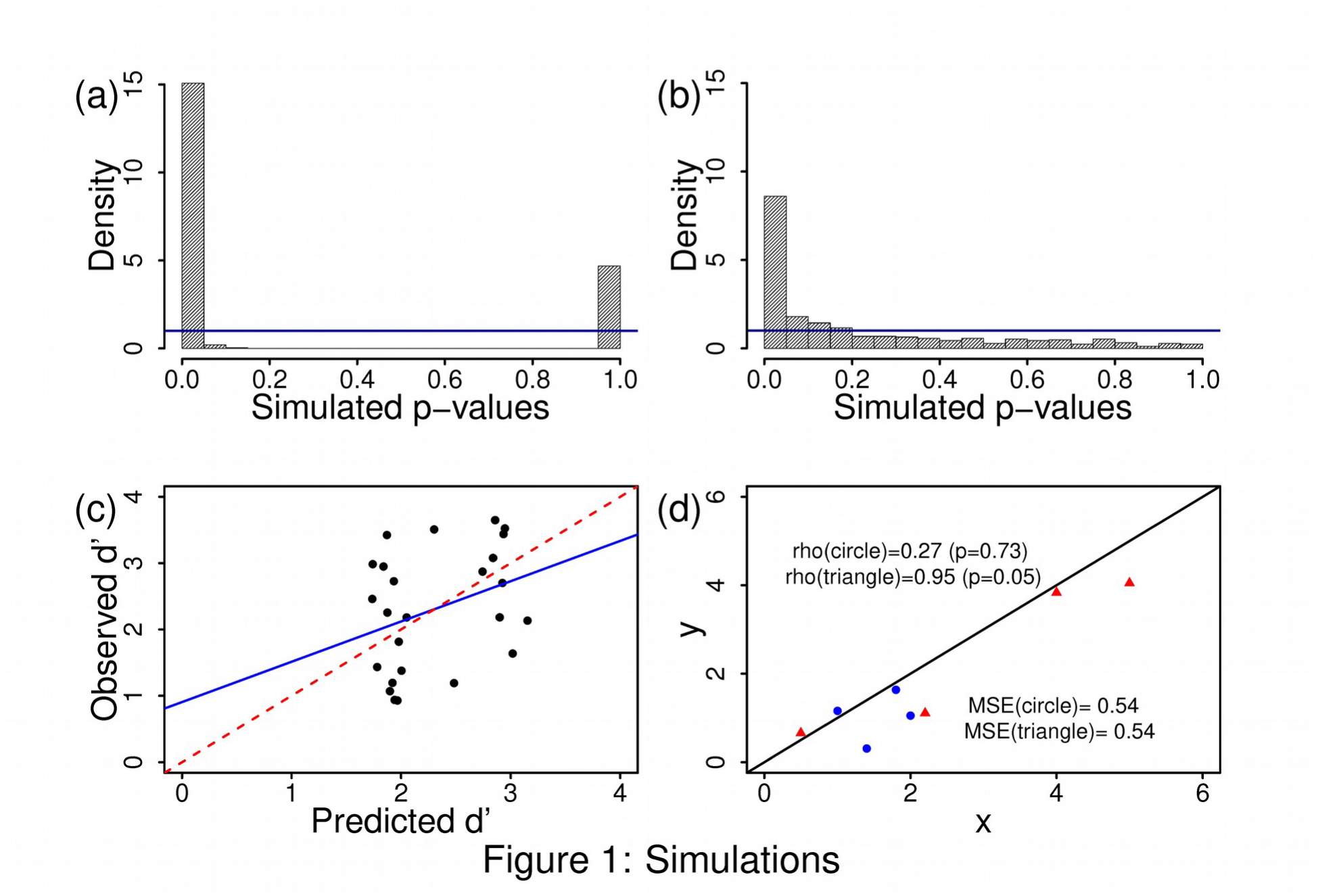
Simulations. (a) Histogram of p-values from 500 simulations of the network validation analysis. The blue line represents the true uniform distribution, which is expected if analyses were unbiased. (b) Histogram of p-values from 500 simulations of the leave-one-out analysis.The blue line represents the true uniform distribution. (c) Scatter plot of observed against leave one out predictions on data simulated as in (b). The blue line represents the regression, the red is the identity. Note that predictions are biased towards the observed mean. (d) Scatter plot of artificial data to illustrate the connection between correlation and heteroscedasticity. The correlation of the blue circles is 0.27 (p=0.73) while the correlation of the red triangles is 0.95 (p=0.05). However, the two point sets have the same distance to the identity (black line) with common MSE equal to 0.54.

### 3.2 Prediction analysis using leave-one-out cross validation (LOOCV)

Following the network validation test, the authors performed prediction analyses on two fMRI data types – data acquired during task performance and data acquired during rest – using a standard LOOCV procedure (see e.g. Hastie et al., 2009) to avoid overfitting. The models were built separately for each training set in the same manner as for the network validation analysis, and predictions were made for the d´ of the left-out participant. Analyses were performed for the positive and negative network strengths as well as for a general linear model based on the two. All analyses were evaluated by examining the Pearson correlation between observed and predicted d´ values. For the analyses based on task related activity, *r* was between 0.84 and 0.87 whereas *r* was between 0.43 and 0.49 for analyses based on resting state data. A main question is how to interpret these correlation tests used to “assess predictive power” (p. 166).

#### 3.2.1 Correlations and R^2 type statistics

In a simple linear regression of x on y, the coefficient of determination, *r*^2^, defined as the proportion of variance in y explained by x (i.e. one minus the ratio of residual to total variation), is calculable as the squared Pearson correlation of x and y. In a multiple regression, *r*^2^ can be calculated as the squared Pearson correlation between the observed and fitted y’s. However, as will be exemplified below, this no longer holds when the fitted values are obtained by cross validation and one generally obtains two different measures by defining an *r*^2^-type measure as the proportion of explained variance (using a cross validated version of residual and total variation) contra the squared Pearson correlation between observed and fitted. The latter definition is taken by the authors (p. 166).

This approach of calculating and testing correlation between observed and LOO predictions entails some methodological pitfalls. Consider a simple example where there are no features available, and we estimate from an intercept-only model thus predicting the left out individual by the mean of the n-1 remaining data points. Here, the LOO prediction and left out observed will be independent, but estimating the Pearson correlation from the sample of n pairs yields a perfect minus one (and thus an *r*^2^ of one). This clear discrepancy stems from the fact that the calculation of the Pearson correlation allows for a recalibration of the model, i.e. a new intercept and slope is fitted. The Pearson correlations are consequently very hard to interpret and will usually be over-dimensioned. Some of these points are also discussed in a recent online blog (Schwarzkopf, 2015).

To illustrate this point, we extended the simulation from the network validation analysis to include leave-one-participant-out cross validation and in each case calculated the correlation between observed and predicted d’. A test for no association was performed for each simulation, and the distribution of the resulting p-values is given in Figure 1b. As the data is simulated under the null hypothesis, the p-values should follow the uniform distribution shown as the horizontal, blue line, but as they do not, the approach is biased.

Another and related consideration worth mentioning pertains to the classical bias-variance trade-off of cross validation estimation of errors with LOO constituting one end of the scale being close to unbiased for the true error but with high variation due to a high positive covariation between estimates (owing to very similar training sets) (Hastie et al., 2009; Kohavi, 1995). In general, one would usually chose a middle ground such as five or ten fold cross validation (Varoquaux et al., 2017).

While we agree with the commendable practise of addressing overfitting by resampling, it is unclear how the reported statistic is to be interpreted, and as shown by simulations, the corresponding test is biased. For this reason, we are in doubt what can be concluded on the basis of the LOOCV test reported by Rosenberg et al. – it is difficult to evaluate whether the findings are robust and whether the predictive power is large enough to support a claim of broad applicability.

### 3.3 Model Selection and Complexity

The authors perform the following analyses to justify the applied level of model complexity. First, two tests were performed with the goal of examining whether d´ was “more closely related to the strength of the whole network than to the strength of individual edges”. They compared the strength of the correlation between d´ and the network strength measure previously obtained to the strongest correlation between d´ and a single edge. For the positive network, the test was not significant (p = 0.2) whereas it was for the negative network (p = 2.4 × 10^−6^), which led the authors to conclude that the network strength captured individual d´ variability better than any single edge. The main purpose of this test thus seems to be whether the selection threshold of p = 0.01 provides better results when applied to the data set than the much more conservative threshold of using the best single edge. Secondly, they test what they call “lesioned” matrices, models excluding one of eight canonical networks. They generally find that little changes in the explanatory power of the model although in multiple cases, the correlation coefficients for the simpler models are in fact higher than for the full model. Surprisingly, at the end of this section the authors caution against simpler models due to the connectivity characteristics of the most important nodes.

The motivation for the model selection is not addressed by the authors. Rosenberg et al. used multiple univariate tests for no correlation between connectivity parameters and the dependent measure with an a priori threshold as a choice of feature exclusion and weighted the remaining features equally when calculating the summary statistic to be used in modelling. This approach seems a bit ad hoc as the applied Steiger’s z test will not take into account the variability stemming from calculating and ranking correlations from the edges. We note that other methods for data reduction exist (e.g. various component or association analyses), and these can be applied to the independent variables alone and thus provide a more principled approach. Supervised approaches that could have been applied include shrinkage regression or other penalisation methods, where one estimates the regularisation parameters (Hastie et al., 2009). When simple, off-the-cuff analyses are employed, it might be of interest to try to assert the sensitivity of the model to subsidiary specifications (e.g. varying the threshold for feature selection). This becomes increasingly true when the model is presented as one representing a whole-brain network. Additionally, we would argue that it is suboptimal that the motivation for choosing a specific model complexity is excluded from the loop when performing cross validation for internal validation.

A main focus of Rosenberg et al. is on the predictive capacity of *complex brain network models*, and although the authors do not claim that their identified model is optimal, they caution against oversimplifying predictive networks to a handful of regions (p. 168) or single features (p. 165), but based on the reported tests, one might just as well have cautioned against making assumptions about whole brain networks. We believe that simple and full models could be compared more systematically. A more principled approach is varying complexity while penalizing the number of variables as more explanatory features cannot reduce explanatory power on the same data. Often one combines such procedures with resampling techniques such as bootstrapping or cross validation to compensate for overfitting.

Based on the reported tests, it cannot be concluded that all parts of the reported network are related to attention (in fact, it appears this is not the case). The external validation tests discussed below do not resolve this issue.

## 4. Permutation test

As mentioned above, the LOO estimates are dependent owing to similar training sets. This is noted by the authors in the last section of the Online Methods, and a permutation test for the association between task performance and edge connectivity had been performed to address the issue. In such an approach, one shuffles the response between individuals and repeats the prediction procedure thus supplying a correlation expected to belong to the null distribution of no association. Repeating this gives an idea of the null distribution of the correlation and the quantile of the observed correlation in this distribution can be evaluated and interpreted as a p-value. This nonparametric permutation distribution is a more correct way to handle the effect of dependence and bias, and therefore we believe that the corresponding p-values obtained in this analysis are more appropriate than the ones reported in the Results section, albeit we retain some scepticism as it is still difficult to interpret LOO correlations that form the basis for the permutations as discussed in section 3.2.

We take the result of this analysis to be a strong indication that connectivity in both the task and rest fMRI datasets are associated with performance in the behavioural task, but we are still not able to evaluate how predictive, and whether it is predictive enough to support the claim of broad applicability. While the permutation test will address the bias stemming from the inflated correlations, it is still a difficult to interpret statistic that is being validated, and the approach does not offer an estimate of the magnitude with which the estimate should be shrunk. Rosenberg et al. could nevertheless provide an idea of the shrinkage factor by supplying a plot of the permutation null distribution.

## 5. External validation

Rosenberg et al. place a large focus on testing the SAN model on external data sets. While testing a model on an external data set generally provides the most reliable measure of its performance, it also entails a substantial interpretational effort when the test data is sampled from a population different from the one on which the model was built. The external validation was performed on resting state data collected at Peking University consisting of 113 children and adolescents, some diagnosed with ADHD, some healthy controls, and the aim was to predict AHDH symptom severity. There is a substantial difference in multiple data attributes, e.g. the examined population, and as a consequence we cannot without extrapolation interpret the correlation as an estimate of the correlation mentioned in the internal data set. As such, the external data correlation does not provide a shrunken estimate.

Another important point to keep in mind is that when performing external validation, statistical significance is dependent on the size of the validation sample, i.e. any nonzero effect will be found in a sufficiently large validation set. Therefore it is essential also in external validation to evaluate the quality of the prediction.

The external validation performed in Rosenberg et al. shows a statistically significant correlation between the SA network and ADHD symptoms and further strengthens the conclusion that (parts of) the network and attention are associated. We return to the results of the analysis below in section 7, where we discuss how one might measure the quality of predictions.

## 6. On neuromarkers

The term neuromarker is related to the more common medical term biomarker of disease, which can be thought of as “a characteristic that can be objectively measured and evaluated as an indicator of normal biological processes, pathogenetic processes, or pharmacologic responses to therapeutic intervention” (Kropotov, 2016, pp. xxiii-xxiv). It was possibly first defined by Gordon (2007) as essentially any objective neuroimaging index of brain structure or function as well as behavioural cognitive tests predictive of some type of disease, but has recently been used more broadly as a neural indicator of a cognitive process.

It is somewhat unclear if Rosenberg et al. hold a similar definition of the term neuromarker as they do not define it. Below, we evaluate the neuromarker proposed by Rosenberg et al. using classic criteria and by examining modelling choices and outcomes from a more traditional statistical perspective on predictive modelling.

## 7. Measures of Predictive Power

Rosenberg et al. place a main focus on predictive modelling and taking the first step towards a brain-based measure of attention (p. 165, p. 170), yet model evaluation is done using only correlation coefficients, an approach more suited to models of association than prediction. Figure 1c is a plot of the predicted and simulated d´s from one of the LOO simulations mentioned above, where the blue line is the regression while the red is the identity (i.e. observed d´ = predicted d´) meant to exemplify that correlation does not signify predictive accuracy. We observe that the predictions fall within a much narrower set of values than those observed, illustrating another problem of the procedure: even though a high correlation between predicted and observed d´ is found, predictions may be biased toward the sample mean. We note that this phenomenon is also observed in Figure 1 of Rosenberg et al. The issue appears more prominent for predictions based on resting state data for which the model predicts that all participants perform within a narrow intermediate range, and, as far as we can see, that children and adolescents diagnosed with ADHD should perform as well as or better than healthy university students. When models are created on the ADHD sample (which contains children/adolescents with/without an ADHD diagnosis) and tested on university students, the model appears to predict that the students have ADHD symptoms comparable to the children/adolescents diagnosed with ADHD. It thus appears that the proposed neuromarker forms the basis of counterintuitive and/or inaccurate predictions. To us, this speaks against the claim of the immediate generalisability of the model, as one cannot meaningfully interpret a single new prediction but must first collect and predict a sizeable sample from the new population to allow the model to recalibrate.

It is unclear why a measure of predictive power should be dependent on the similarity of the predictions as indeed the correlation is. Figure 1d shows two sets of points around the identity line, where the identity corresponds to perfect agreement. One sees that the set of blue circles have a lower correlation than the red triangles, the difference transcending that of statistical significance on a 5% level. However, the two sets have the exact same distance to the identity, the red being simply more heterogeneous. Another related property of the correlation that makes it unattractive as a predictive accuracy measure is that it is invariant to scaling of the axes. These properties, however, make the correlation an appropriate measure of discrimination, i.e. how easily it is to separate or rank the population based on the model. It is therefore also natural that the correlation depends on the variance in the population since it is easier to discriminate in a heterogeneous population.

An often used measure of predictive accuracy for continuous data is the mean square error (MSE), defined to be the expected squared deviation from observed to predicted. This has the advantage of often being relatable to the width of an asymptotic prediction interval and therefore giving a quantity that can be interpreted lucidly on the actual scale of the observations. Other measures include the Gini index and calibration curves, measures also relevant for discrete observations (Harrell, 2015). We believe such measures would be practically useful additions to the reported correlations.

Given that Rosenberg et al. propose a broadly applicable neuromarker and claim specifically that the “SAN model can also predict symptoms of attention deficit hyperactivity disorder” (p. 165), they could additionally have supplemented the analysis by evaluation of predictive accuracy using standard methods for neuromarkers – such as sensitivity/specificity – instead of using only a method for evaluating predictive models in general. For example, Kropotov (2016, p. xxiii) mentions that an ideal neuromarker for ADHD should have a diagnostic sensitivity (probability of correct positives) of at least 80% and a diagnostic specificity (probability of correct negatives) of at least 80%.

Sensitivity and specificity depends on a selected threshold (cut-off) value and are therefore typically reported for a range of thresholds. Such thresholds can be set for Rosenberg et al.’s data predicting ADHD using the GLM data presented in Figure 3: a predicted d´ threshold is selected, and the number of ADHD (hits) and non-ADHD (false alarms) children below this threshold can be counted, and from these, sensitivity and specificity can be calculated. Using this method, we found that a threshold achieving a specificity of around 80% results in a sensitivity of around 40-45%. Similarly, to achieve a sensitivity of 80%, specificity is around 40-45%. With a more balanced threshold, we are able to achieve both sensitivity and specificity of around 60%. Thus, if we apply a standard method for neuromarker evaluation in the context of ADHD, the method of Rosenberg et al. does not appear to fulfil the sensitivity/specificity requirements and is only slightly better than random guessing.

Taken together, we thus believe that the model evaluation method of using *r*^2^ between the predicted and observed values have a number of disadvantages. As already pointed out, such measures are difficult to interpret when predictions are obtained by LOOCV. For predictions on healthy participants, the analyses could have been supplemented with accuracy measures such as the mean squared error. For predictions of ADHD symptoms, analyses could have been supplemented with more traditional evaluation measures for neuromarkers such as sensitivity and specificity. For both alternative evaluation methods, the results appear less impressive.

## 8. Concluding discussion

In cognitive neuroscience, whether we define a neuromarker as a measurable neural indicator of a cognitive characteristic or disease, we would expect that neuromarker to be a precise and reliable surrogate. In their article, Rosenberg et al. (2016) claim that “whole-brain functional network strength provides a broadly applicable neuromarker of sustained attention” (p. 165). While we do not doubt that the network has some predictive value for attention, we have presented three concerns related to the claim which we recount in the following adding a few recommendations for future studies using similar methods.

First, we argued that two of the most central analyses, the network validation test and the LOOCV test are performed in a biased and difficult to interpret manner, and the analyses on the collected data therefore amount to a nonparametric association test (the permutation test) with no working estimate of the effect. In regards to the specific estimation of an *r*^2^ type statistic from cross validation we would recommend using the CV version of residual and total error in its calculation. We further argued that the correlations resulting from external validation cannot be taken as estimates in the internal data due to substantial differences in data and populations. For the same reason it is not clear how the correlations should be interpreted.

Second, we are concerned that a focus on “whole brain” connectivity could be misleading as model comparisons were not performed ideally, and the analyses that were conducted indicated that smaller networks were equally or more predictive. As a general recommendation, data reduction is usually unproblematic, at least in terms of overfitting, when it is done in an unsupervised manner, i.e. blinded to the response. Supervised learning where one selects features using the response can be performed for example by maximising some information criterion that should penalise for model complexity. It is important to account for the resulting overfit when reporting model performance. As a final suggestion we propose that when a neuromarker of potential interest to classification is reported, its sensitivity and specificity (or similar measures) are part of the report as these could be of use to clinicians working with biomarkers. Proposing a neuromarker as broadly applicable could be misleading without an estimate of its accuracy.

Third, we argued that a narrow focus on difficult-to-interpret, relative predictive capabilities of the model makes it even more difficult to evaluate the predictive value of the model. As in clinical research in general, the methodological focus on significance testing, here whether the network significantly correlates with attention, is often of small interest. Rather the focus should be on the effect size and the certainty with which it was determined. While it is not obvious how we should measure an effect in predictive modelling, we have tried to argue that it should be constituted by more than a correlation. We also argue that estimates should be chosen and shrunk to reflect a realistic performance of the model on a new sample, another point where we find the applied approach suboptimal. Claims of the network strength being a “robust” and “broadly applicable” predictor are consequently very difficult to verify from the article, and applicability outside the original dataset appears dependent on model recalibration. Furthermore, the authors do not specify their criteria for claiming broad applicability, thus making the claim even harder to evaluate.

When summarising a model, it is advisable to include multiple performance measures to describe the performance. These measures will usually be too optimistically estimated which can be sought remedied by resampling for example by enhanced bootstrap (Efron, 1983; Harrell, 2015). Shrinkage can alternatively be built into the estimation in regularised regression, which involves tuning of regularisation parameter(s) and considerations on data scaling.

In summary, we find that the article by Rosenberg et al. provides ample proof that parts of the SAN network is associated with capacity for sustained attention through permutation tests and external validation. We argue, however, that the performed analyses do not warrant the conclusion that the network is a robust, broadly applicable, predictive neuromarker.

## Acknowledgements

We thank Bo Martin Bibby for comments on the manuscript.

